# Effects of substrate stiffness on the biological behavior of human umbilical vein endothelial cells

**DOI:** 10.1101/803312

**Authors:** Hua Pei, Liang Li, Kejun Liu, Wenming Wang, Bowen Jiang, Yiji Li, Li Zuo

## Abstract

**Background:** The biophysical attributes of a substrate can directly influence endothelial cell behaviors. Here, we explored substrate stiffness and its biological impact on human umbilical vein endothelial cell (HUVEC) behavior, representing different anatomical sites and differentiation states *in vivo*.

**Material and Methods:** HUVECs were cultured on both stiff substrate (25 kPa hydrogel GEL) and tissue culture plastic (TCP). Cell cytoskeleton and proliferation were detected by immunofluorescence and BrdU assays, respectively. The protein and gene expression levels of connexin 40 (CX40) were ascertained via Western blotting analysis and quantitative real-time polymerase chain reaction. Glycosaminoglycan (GAG) content was determined using a sulfated GAG detection kit.

**Results:** Results showed that actin stress fiber density and HUVEC proliferation both decreased, whereas CX40 expression and GAG content both increased in the cells grown on the stiff substrate compared to cells grown on the TCP.

**Conclusions:** Following culture on the stiff substrate, the biological behavior of the HUVECs differed obviously to those cultured on the TCP. Our results suggest that the state of the cells cultured on the stiff substrate may be similar to their phenotypic state *in vivo*.

## BACKGROUND

Vascular endothelial cells (VECs) are a monolayer of flat or polygonal cells that cover the surface of the vascular endothelium and maintain cardiovascular system homeostasis in various ways [1]. *In vitro* culture of VECs is an important tool for studying pathogenesis. However, current *in vitro* culture models remain unsatisfactory, particularly in regard to the large differences in the cell environment. Normal VEC wall stiffness ranges from 3 to 70 kPa, and even after atherosclerosis, stiffness is no more than 200 kPa [2-4]. Conversely, the stiffness of universal tissue culture plastic (TCP) falls within the 2–4 GPa range, far higher than that of VECs found within the human body under physiological and pathological states.

The stiffness or rigidity of a culture substrate can markedly affect the biological behavior of cells. For example, changes in matrix stiffness are closely related to the tumorigenicity of tumor regenerated cells [5, 6]. Furthermore, mammalian early embryonic cell clusters can only differentiate into the three germ layers *in vitro* under appropriate base stiffness [7]. Differences in matrix stiffness in macrophages can also affect cell phagocytosis and cytokine secretion [8]. In the case of VECs, the elastic stiffness of the vascular basement membrane and underlying matrix can affect important biological functions, such as survival, replication, migration, and steady state maintenance [9]. VECs undergo pathological changes with increasing substrate stiffness, which impacts the balance of matrix metalloproteinases and increases the deposition and cross-linking of matrix collagen, thus accelerating the hardening of the cardiovascular system *in vivo* [10-13]. Therefore, in the present study, *in vitro* VECs were investigated under different substrate stiffnesses.

Using current *in vitro* culture models to study the physiological and pathological functions of VECs is not ideal due to the large differences in substrate stiffness *in vivo* and *in vitro*. As such, we cultured human umbilical vein endothelial cells (HUVECs) on hydrogels (GEL) with a substrate rigidity close to actual physiological vascular stiffness (25 kPa). As a control, we also used TCP to evaluate changes in HUVECs cultured in both systems (i.e., modifications in morphology, proliferation, and connexin 40 (CX40) and GAG expression levels). We evaluated whether *in vitro* culture of the HUVECs could be improved by increasing substrate elasticity.

## MATERIAL AND METHODS

All research protocols involved in the current study were approved by the Medical and Health Research Ethics Committee of Hainan Medical University in Haikou, China. Furthermore, all patient procedures followed the standards set by the Helsinki Convention [14], with written informed consent provided by all relevant patients and participants.

### Experiment materials

The reagents used in this experiment included endothelial growth factor (EGF) (Becton, Dickinson and Company, USA) and trypsase (Thermo Scientific, USA).

### HUVEC culture

The HUVECs were incubated at 37 °C under 5% CO_2_ in an appropriate medium (1% VEGF-M199), with the addition of growth factor (1% M199, Thermo Scientific, USA) and fetal bovine serum (10%, FBS, Biological Industries, USA). After 3–4 generations of resuscitation, cells in good condition were used for the subsequent experiments.

### Cell area and vertical and horizontal axis analysis

The HUVECs were inoculated in 25 kPa 6-well GEL plates (Softwell, Matrigen Life Technologies, USA) or 6-well TCP plates at a density of 1 × 10^4^ cells/ml for 48 h, then fixed, treated, and stained with 4% paraformaldehyde, 0.5% Triton X-100, and FITC phalloidin (0.5 µg/ml, Sigma-Aldrich LLC, USA), respectively, and finally observed under an inverted fluorescence microscope (Olympus, Japan). Various parameters (i.e., longitudinal axis, transverse axis, projection area) of the cells grown on the two different culture substrates were measured using ImageJ. Means were calculated and used to determine the effects of substrate stiffness on cell proliferation and growth. At least 100 cells grown on each substrate were selected for measurement, with three replicates established. All measurements were pooled and averaged.

### BromodeoxyUridine (5-Bromo-2-DeoxyUridine, BrdU) proliferation test

To HUVECs were incubated at 37 °C under 5% CO_2_ in 6-well GEL and TCP plates at a density of 1 × 10^5^ cells/ml in appropriate culture medium (M199) supplemented with BrdU reagent (Sigma-Aldrich LLC, USA). After 48 h, the cells were fixed with 4% paraformaldehyde (30 min) and rinsed with PBS buffer, with denaturation and neutralization initiated by the addition of 2 M HCl and sodium borate buffer (15 min each), respectively. The cells were subsequently blocked (1 h) with 10% BSA, supplemented with anti-BrdU primary antibody, thrice washed with PBS, then supplemented with FITC-labeled secondary antibody and DAPI-containing tablets. The cells were finally observed using 20 visual fields at 100× magnification with an inverted fluorescence microscope (Olympus, Japan). Cells were counted using ImageJ software and HUVEC proliferation was calculated. All experiments were repeated independently in triplicate, from which means were then calculated.

### CX40 mRNA expression

We used a total RNA kit (Tiangen Biochemical Technology Co., Ltd., China) to isolate RNA from the HUVECs cultured on the two different substrates (i.e., GEL and TCP). Quantitative real-time polymerase chain reaction (qRT-PCR) was undertaken using TaqMan One-Step RT-PCR Master Mix (Applied Biosystems, USA). The cDNA was synthesized at 42 °C for 60 min and at 70 °C for 5 min. The qRT-PCR conditions were: 10 min at 95 °C, 15 s at 95 °C, and 60 s at 60 °C for 40 cycles; followed by 15 s at 95 °C, 1 min at 60 °C, and 15 s at 95 °C for melting curve analysis. The primers used here, including human CX40 forward primer: GGGCACTCTGCTCAACACCT and reverse primer: TGAAGCCCACCTCCATGGT, were developed by Beijing Liuhe Huada Gene Technology Co., Ltd. (China). Data were normalized to glyceraldehyde-3-phosphate dehydrogenase (GAPDH), with forward primer: GCACCGTCAAGGCTGAGAAC and reverse primer: TGGTGAAGACGCCAGTGGA. The relative expression of CX40 mRNA was calculated.

### CX40 protein expression by Western blot analysis

Western blotting was carried out as described previously [15]. The HUVECs were cultured in 6-well GEL and TCP plates at a density of 3 × 10^5^ cells/ml and then collected at 48 h. Protein was lysed by SDS and quantified via bicinchoninic acid assay (BCA). A 10% polyacrylamide gel was prepared, with 25 μg loaded into each well. Proteins were electroblotted onto PVDF membranes, followed by overnight (4 °C) incubation with anti-CX40 primary antibody (1:1□000 dilution) (Abcam), 1-h (room temperature) incubation at with horseradish peroxidase-labeled secondary antibody (Abcam) (1:2□000 dilution), and gel imager analysis.

### Glycosaminoglycan (GAG) content detection

We determined GAG content using a Blyscan Sulfated Glycosaminoglycan Assay (Bicolor, UK) as per the detailed protocols provided by the manufacturer. The HUVECs were incubated in 6-well GEL and TCP plates at a density of 3 × 10^5^ cells/ml. After 48 h, the cells were thrice rinsed with PBS and then mixed with PBS solution containing 125 ng/ml papain. After 24-h (60 °C) incubation in a shaker and 5-min centrifugation at 10□000 rpm, the resulting supernatant was harvested and stained with methylene blue dye. GAG content was then measured at a wavelength of 656 nm with a spectrophotometer (Bio-Rad SmartSpec Plus, USA).

### Statistical analysis

All data analyses were performed using SPSS 13.0. Comparisons for cell area, vertical and horizontal axes, CX40 expression, and GAG content were evaluated via *t*-tests, with *P* < 0.05 deemed a measure of statistical significance.

## RESULTS

### Changes in HUVEC area and vertical and horizontal axes based on substrate stiffness

Based on inverted fluorescence microscopy, most HUVECs cultured on TCP were found to be elongated (Figure 1), with an increased cell surface area (*t-*test; *P* < 0.05) (Table 1). In comparison, HUVECs cultured on GEL were rounded, with significantly shorter long and short axes (Figure 1) (*t-*test; both *P* < 0.05) (Table 1).

**Table 1.**
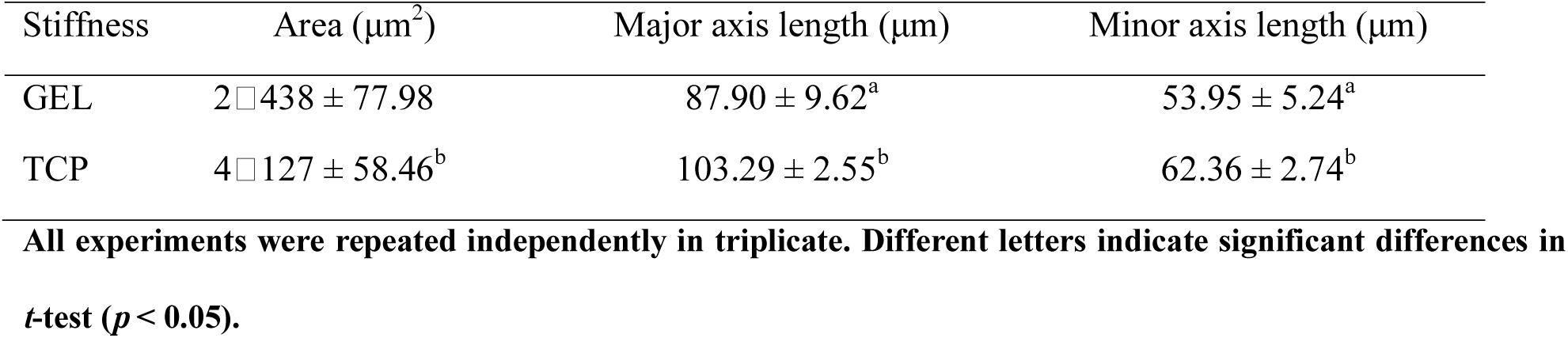
Surface area and length of vertical and horizontal axes of HUVECs cultured under different substrate stiffness.

**Figure 1.**
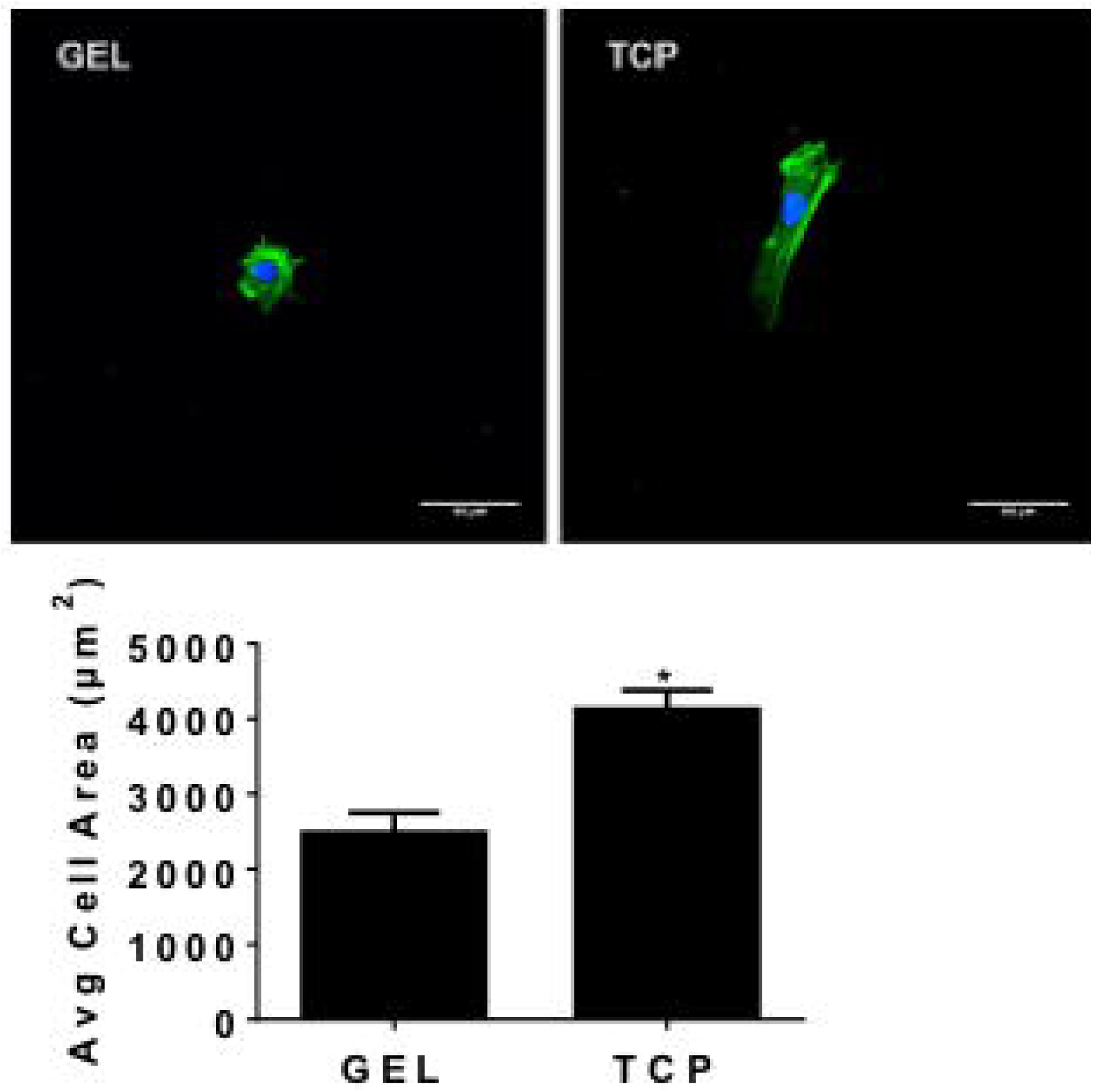
HUVEC staining with FITC phalloidin. All experiments were repeated independently in triplicate. Different letters indicate significant differences in *t-*test (*p* < 0.05).

### Changes in HUVEC proliferation based on substrate stiffness

Based on the BrdU test, the proliferation of HUVECs cultured on TCP was 9.01 ± 0.14%, whereas the proliferation of HUVECs cultured on GEL was only 1.47 ± 0.32% (*t-*test; *P* < 0.05) (Figure 2).

**Figure 2.**
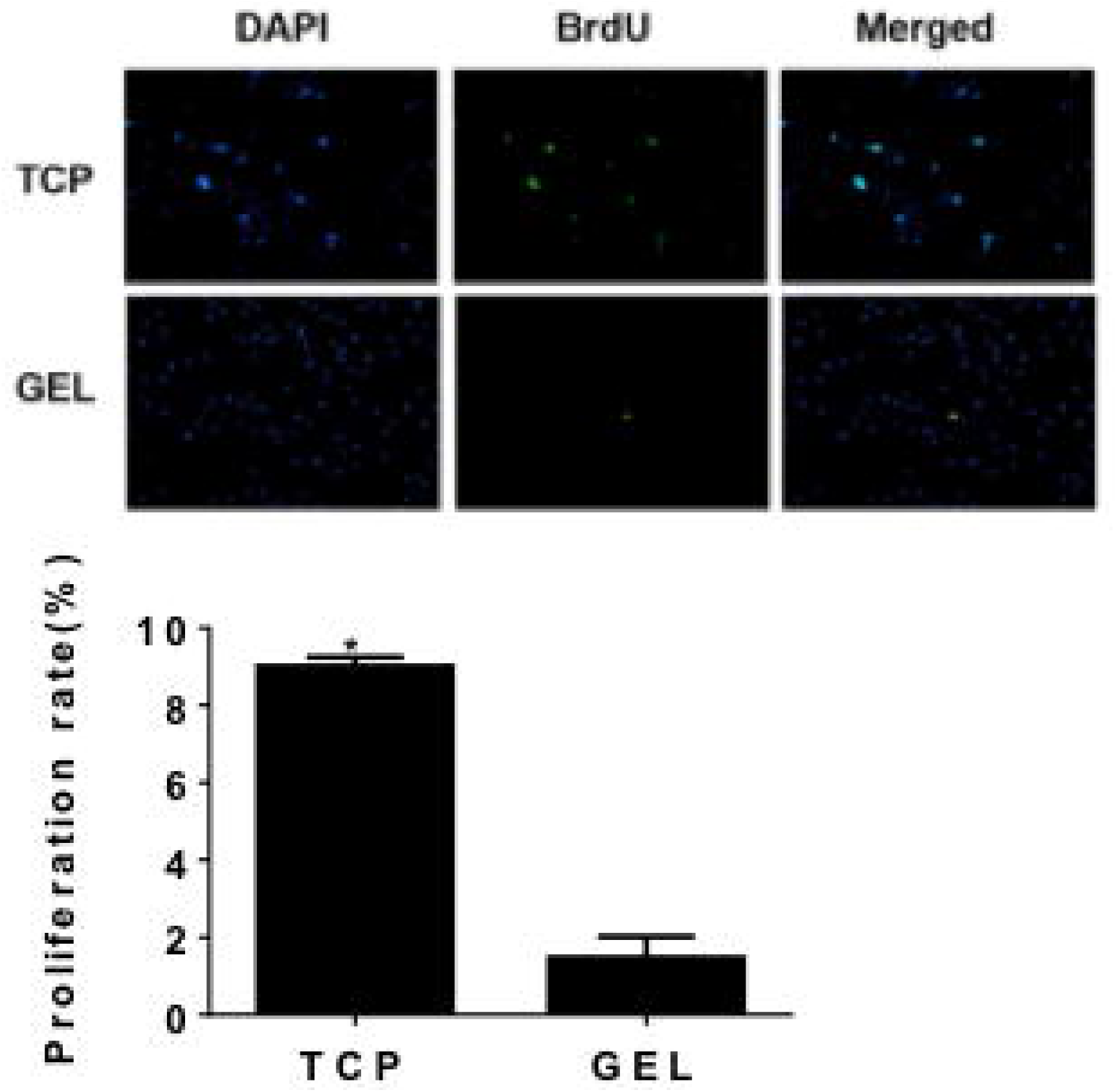
Effects of substrate stiffness/rigidity on proliferation of HUVECs based on bromodeoxyuridine (BrdU) assay. All experiments were repeated independently in triplicate. Different letters indicate significant differences in *t-*test (*p* < 0.05).

### Changes in CX40 mRNA and protein expression based on substrate stiffness

Both the CX40 mRNA and protein expression levels were significantly higher in cells cultured on GEL than those cultured on TCP (*t-*test; *P* < 0.05, Figure 3 and Figure 4, respectively).

**Figure 3.**
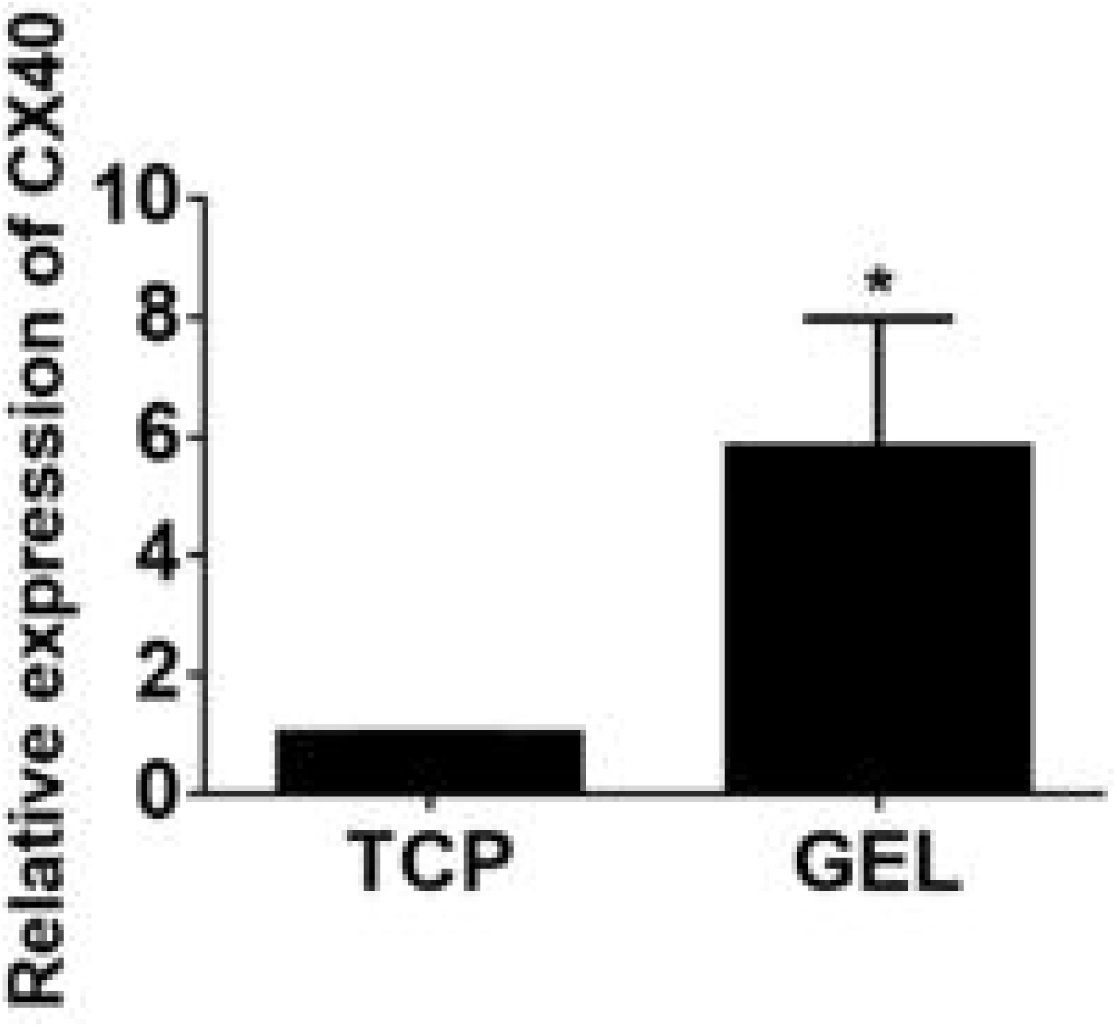
Effects of substrate stiffness/rigidity on CX40 mRNA expression in HUVECs. All experiments were repeated independently in triplicate. Bars are standard deviation (SD) based on three replicate samples. Different letters indicate significant differences in *t-*test (*p* < 0.05).

**Figure 4.**
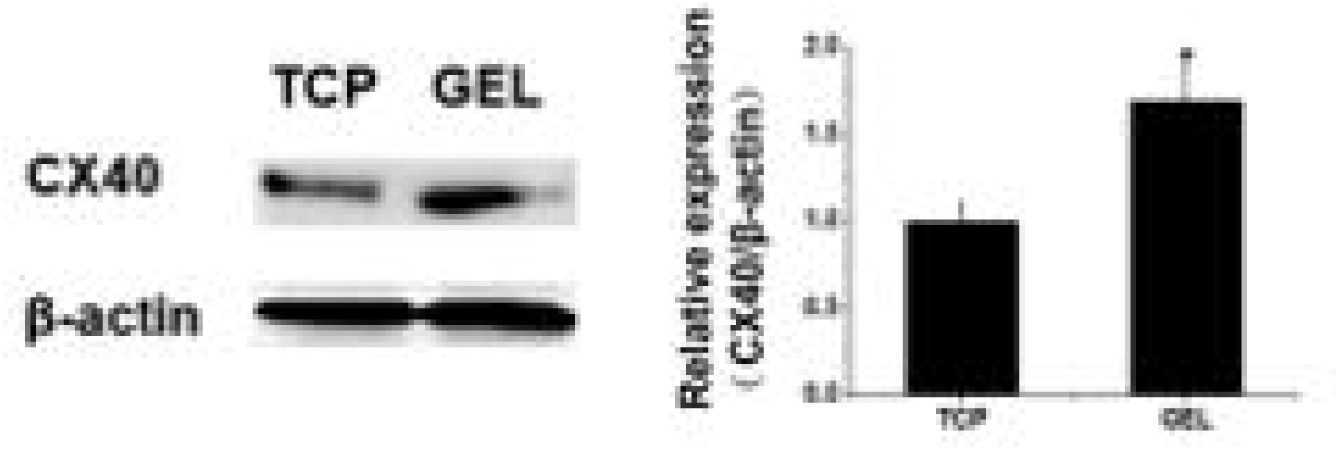
Effects of different substrate stiffness/rigidity on CX40 protein expression in HUVECs. All experiments were repeated independently in triplicate. Bars are standard deviation (SD) based on three replicate samples. Different letters indicate significant differences in *t-*test (*p* < 0.05).

### Changes in GAG content based on substrate stiffness

GAG content was significantly higher in HUVECs cultured on GEL than those cultured on TCP (*t-*test; *P* < 0.05, Figure 5).

**Figure 5.**
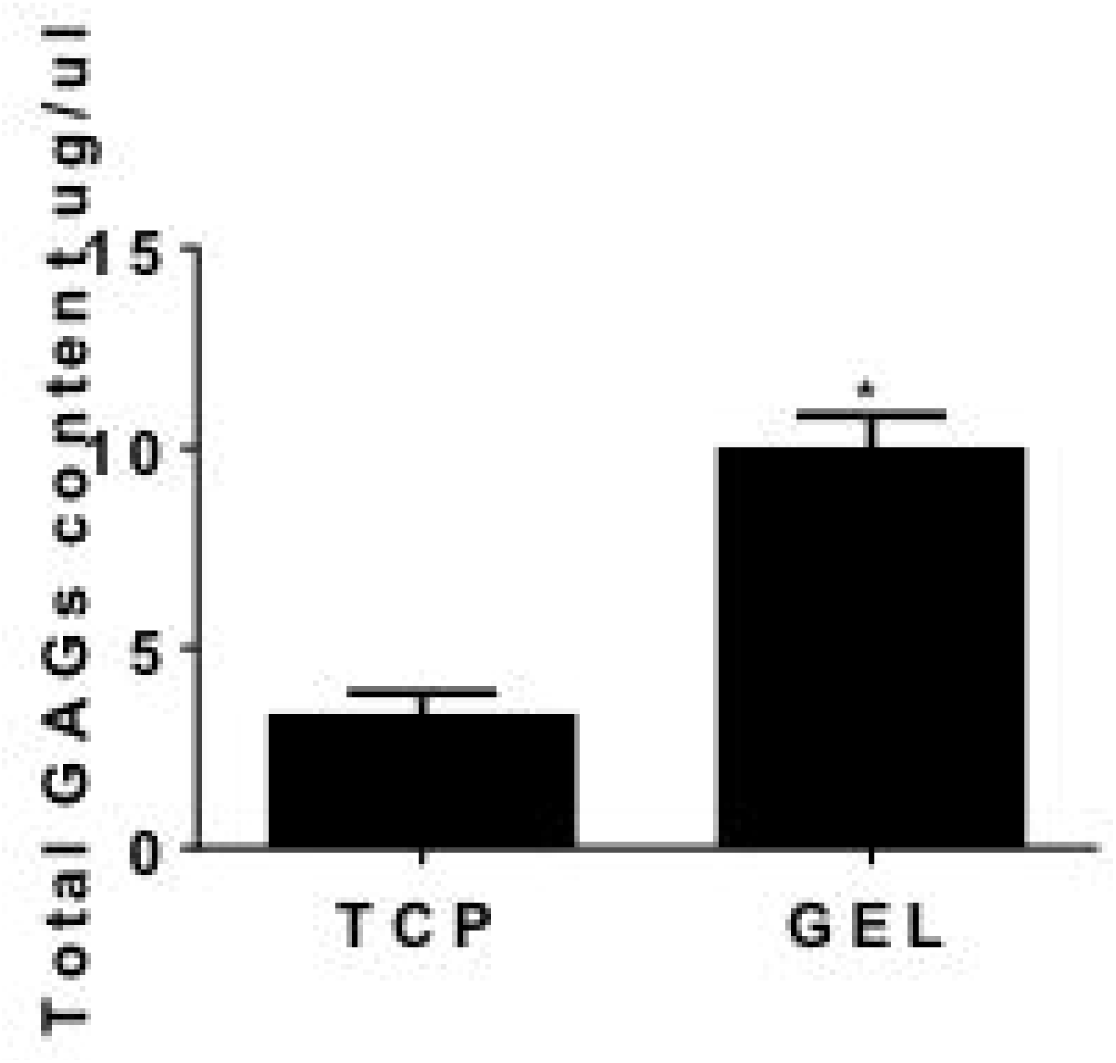
Total GAG content in HUVECs cultured under different substrate stiffness/rigidity. All experiments were repeated independently in triplicate. Bars are standard deviation (SD) based on three replicate samples. Different letters indicate significant differences in *t-*test (*p* < 0.05).

## DISCUSSION

Although most VECs are found at rest, when they are damaged or stimulated by inflammation they are transformed into a proliferative migratory state or activated state, respectively. VECs cultured using traditional *in vitro* methods are similar to *in vivo* proliferative migratory cells [16, 17]. Here, we showed that HUVECs cultured on GEL had a smaller area and shorter vertical and horizontal axes than cells cultured on TCP. According to previously reported results [18] and the cell morphology and proliferation rate of the current study, our GEL-cultured HUVECs were more similar to VECs in resting state than the TCP-cultured cells.

On the surface of VECs, GAG is mainly composed of proteoglycans and membrane glycoproteins [19]. GAG is necessary to maintain the normal structure and function of VECs and participates in the selective permeability barrier formation of the vessel wall and interaction between blood cells and VECs [20]. GAG content in VECs can significantly decrease due to high blood pressure, atherosclerosis, and other cardiovascular and cerebrovascular diseases [21]. Challengingly, the expression of cell surface GAG is markedly decreased during traditional *in vitro* culture of VECs [22]. In our study, however, the overall content of GAG was significantly up-regulated in the HUVECs cultured on GEL compared to cells cultured on TCP.

As an important gap junction gene, CX40 plays an integral role in the formation of these junctions, via which essential intercellular energy and information are exchanged between adjacent cells [23]. However, CX40 expression can decrease in certain diseases, such as atherosclerosis, diabetes mellitus, and pulmonary hypertension [24, 25]. Previous research has reported that CX40 expression in VECs following *in vitro* culture exhibits a downward trend [26]. In this study, however, CX40 expression in the HUVECs cultured on GEL was higher than that in cells cultured on TCP.

## CONCLUSIONS

The HUVEC culture model was optimized *in vitro* when using a hydrogel with near physiological base elasticity, and is thus a more suitable model for research work. However, VECs demonstrate a high degree of heterogeneity *in vivo*, and different sized blood vessels can exhibit differences in base stiffness [27]. Therefore, we recommend that the optimum substrate stiffness of VECs derived from different tissues should be evaluated *in vitro*.

## Authors’ contributions

HP and LZ conceived the study and coordinated its implementation. HP, LL, and LZ participated in the experimental design. HP, LL, KL, WW, BJ, and YL performed the experiments and drafted the manuscript, which was critically revised by HP and LZ. All authors read and approved the final version of the manuscript.

## Acknowledgements

We are very grateful to any anonymous reviewers for their helpful comments, which have significantly improved the quality of our paper.

